# Low-Latency sEMG Conditioning with Frequency-Tracked Harmonic PLI Cancellation and Exponential RMS Envelope Estimation

**DOI:** 10.64898/2026.06.11.724714

**Authors:** Hong U Lo, Zibo Gao, Hoi Fong Loi, Sok Kin Cheng

## Abstract

Surface electromyography (sEMG) interfaces require interference-resistant acquisition, faithful preprocessing, and a responsive envelope for downstream control. This work presents a reproducible, analog-aware digital conditioning pipeline comprising an eighth-order Butterworth band-pass, a narrow static notch, a recursive-oscillator two-harmonic normalised least-mean-squares (NLMS) canceller, a rolling mains-frequency tracker, and an exponentially weighted RMS (EMRMS) envelope. The analog common-mode analysis is separated into a complex frequency-domain decomposition and a scalar CMRR sensitivity screen; the latter is explicitly treated as a pre-prototype approximation rather than a substitute for circuit simulation or bench measurement. A running-sum rectangular RMS requires *O*(1) arithmetic and *O*(*L*) memory, whereas EMRMS requires *O*(1) arithmetic and one dynamic power state. At *f*_*s*_ = 2 kHz and *τ* = 25 ms, the EMRMS power smoother has a weight-centroid delay of 24.75 ms and a noise-equivalent rectangular length of 100.0 samples. In a controlled drifting-line experiment, the tracker achieved an RMSE of 0.118 Hz; tracked two-harmonic cancellation improved whole-record reconstruction SNR by approximately 19.3 dB relative to the static-notch baseline. In a separate 12-channel synthetic ablation, median reconstruction SNR was 11.5 dB for tracked harmonic cancellation, compared with 8.8 dB for fixed-frequency harmonic cancellation and −11.2 dB for the static-notch baseline. Delay-aligned EMRMS and length-200 running-RMS envelopes had correlation *ρ* = 0.994. The host-Python fast path required a median of 13.0 µs per sample and a 99th percentile of 24.9 µs; accepted frequency updates required a median of 161 µs and should be scheduled as a supervisory task on embedded hardware. All numerical performance results reported here are controlled synthetic experiments. The accompanying real-data workflow never substitutes generated data when recordings are absent; multi-subject and embedded-hardware validation remain future work.

## 1 Introduction

Surface electromyography (sEMG) measures the electrical activity associated with active motor units through electrodes placed on the skin. The recorded waveform depends not only on the underlying physiology but also on electrode placement, electrode-skin impedance, differential acquisition, analog conditioning, sampling, and digital preprocessing. Best-practice guidance therefore recommends that these stages be reported as a connected measurement chain rather than as isolated algorithmic choices (Merletti and Cerone, 2020; Clancy et al., 2023; Besomi et al., 2024). In prosthetic control, rehabilitation robotics, exoskeletons, and human–machine interfaces, this chain must also respond quickly enough to avoid degrading control performance (Farrell and Weir, 2007; Smith et al., 2011).

Two technical problems motivate the present work. First, the differential sEMG signal is small relative to common-mode pickup from the electrical environment. High input impedance, balanced electrode interfaces, instrumentation-amplifier common-mode rejection, and driven-right-leg (DRL) feedback are therefore central design considerations in wearable front ends (Winter and Webster, 1983; Wang et al., 2023; Kledrowetz et al., 2023; Cerone et al., 2019). Second, residual power-line interference (PLI) can remain after analog conditioning. Static notches, comb filters, adaptive cancellers, and time-frequency methods have all been used to attenuate PLI in biopotential measurements (Boyer et al., 2023; Tomasini et al., 2016; Tabakhi et al., 2023). Methods that jointly estimate the fundamental frequency and multiple harmonics are more robust to drift than a fixed-frequency two-weight LMS, but they require additional state and arithmetic (Keshtkaran and Yang, 2014).

Envelope estimation introduces a separate smoothness–responsiveness trade-off. Rectangular RMS, average rectified value, linear envelopes, and adaptive estimators are widely used, with the selected smoothing scale depending on the physiological and control objective (Ranaldi et al., 2018; Nougarou et al., 2023; Clancy et al., 2023). A causal rectangular RMS can be implemented efficiently with a running sum, but it retains a length-*L* history. An exponential power smoother uses one state, making it attractive for memory-constrained deployment. The exponential recursion itself is classical; the contribution here is to account for its delay, memory, startup behaviour, filter interaction, and PLI assumptions within a reproducible system.

The individual filtering, adaptive-cancellation, and exponential-smoothing operations used in this work are established techniques. The contribution lies in their transparent two-rate integration, the explicit separation of mains-frequency estimation from adaptive amplitude–phase cancellation, and the joint accounting of interference suppression, envelope delay, memory, arithmetic cost, and implementation constraints. The software further prevents silent substitution of generated data when a real-data command is requested and records all numerical outputs used by the manuscript.

Accordingly, this study develops a unified framework that connects front-end common-mode rejection, adaptive PLI suppression, envelope estimation, and implementation-level resource analysis. A frequency-domain decomposition of common-mode-to-differential conversion is paired with a clearly labelled scalar sensitivity screen for pre-prototype exploration. Running-sum rectangular RMS and EMRMS are compared through arithmetic complexity, memory, centroid delay, equivalent window length, and startup behaviour. Time-varying interference is addressed by a fixed-rate supervisory tracker with confidence gating and bounded frequency updates, coupled to a recursive-oscillator two-harmonic NLMS canceller. The implemented digital chain is also characterised in terms of realised filter order, arithmetic operations, dynamic state, host execution time, and frequency-dependent group delay.

## 2 Methods

### 2.1 System scope and processing chain

The implemented digital processing chain operates at *f*_*s*_ = 2 kHz. The acquired signal is first passed through an eighth-order Butterworth band-pass filter covering 20–450 Hz, implemented as four second-order sections (SOS), followed by a second-order notch filter centred at 50 Hz with a quality factor of *Q* = 30. Residual power-line interference is then attenuated by a two-harmonic normalised least-mean-squares (NLMS) canceller. Its recursive oscillator may operate at a fixed frequency or be updated periodically by the bounded-rate supervisory frequency tracker. Finally, an exponentially weighted RMS (EMRMS) estimator with a time constant of *τ* = 25 ms produces the control-oriented signal envelope.

The analog component of the present study is restricted to mathematical sensitivity analysis intended to support pre-prototype design. No fabricated analog front end or circuit-level SPICE model is evaluated in this version, and the study does not include injected common-mode bench testing, a fixed-point microcontroller implementation, RTOS deadline and jitter measurements, or direct power-consumption measurements. These elements therefore remain outside the scope of the claims made in the current work.

### 2.2 Analog common-mode rejection

A physically complete front-end model is frequency dependent. Let *V*_cm,b_ denote the body common-mode voltage before DRL action and let *T*_DRL_ (*j*Ω) denote the DRL loop gain. To first order, the residual common-mode voltage presented to the differential input is *V*_cm,in_ = *V*_cm,b_ / [1+ *T*_DRL_ (*j*Ω)] . Let *H*_INA,cm_ (*j*Ω) denote the instrumentation amplifier’s internal common-mode-to-differential conversion relative to this residual input voltage, and let *H*_Δ*Z*_ (*j*Ω) denote the corresponding conversion caused by electrode and input-impedance imbalance. Under these reference definitions, a useful first-order closed-loop decomposition is

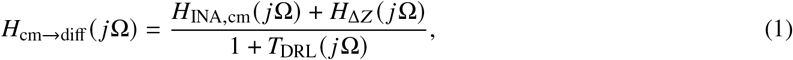

with

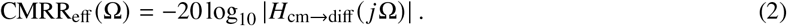

This decomposition preserves complex phase and makes the voltage reference explicit. It remains a first-order representation rather than a circuit derivation: cable capacitance, protection components, complex electrode impedance, loop stability, saturation, and amplifier frequency response must be evaluated using a specified topology.

For preliminary sensitivity screening at one interference frequency, we also use

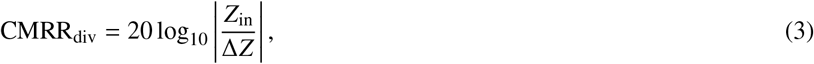

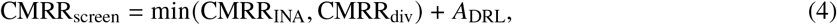

where *A*_DRL_ = 20 log_10_ | 1 +*T*_DRL_| is interpreted as closed-loop common-mode attenuation attributable to the DRL path, not raw op-amp open-loop gain. Equation (4) neglects vector addition and is used only as an upper-bound-style design screen.

### 2.3 Running-sum RMS and EMRMS

A length-*L* causal rectangular RMS is

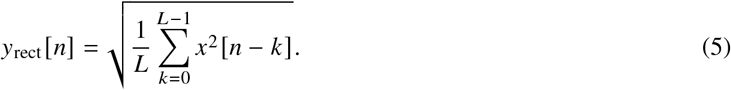

A running sum gives

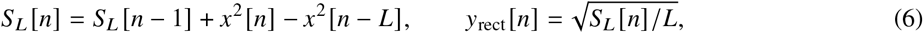

so a practical rectangular RMS uses *O* (1) arithmetic and *O* (*L*) memory. In the reported zero-state implementation, *x* [*n*] = 0 for *n* < 0 and the denominator remains *L* during startup; a partial-window normalisation would define a different transient and is not used here.

The EMRMS recursion is

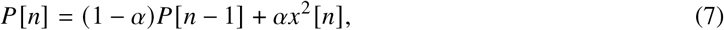

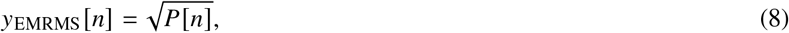

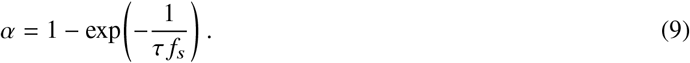

It stores one power state and requires *O*(1) arithmetic and *O*(1) state. The exponential weighting impulse response is *h*[*k*] = *α*(1 − *α*)^*k*^ for *k* ≥ 0. Its weight-centroid delay is

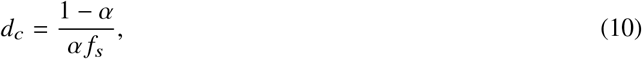

and its noise-equivalent rectangular length is

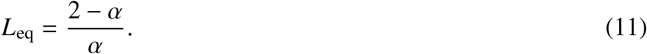

At *f*_*s*_ = 2 kHz and *τ* = 25 ms, *d*_*c*_ = 24.75 ms and *L*_eq_ ≈ 100.0 samples. The implementation additionally provides optional startup bias correction,

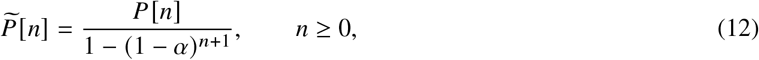

for offline comparison. It is disabled by default because it adds a division and changes the fast-path operation count.

### 2.4 Harmonic normalized-LMS cancellation

For *K* harmonics and a supplied phase *θ* [*n*], the estimated PLI is

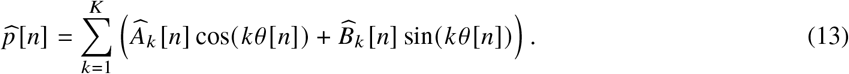

The cleaned output is 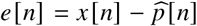. Since each harmonic contributes a cosine–sine pair with unit squared norm, the complete regressor has constant squared norm *K*. The normalized update can therefore be written

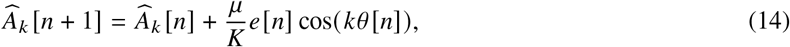

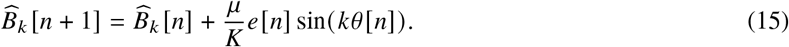

The usual sample-wise normalized-LMS range is 0 < *μ* < 2 (Haykin, 2008). The default implementation uses *K* = 2 and *μ* = 0.02. A leakage term is available but disabled in the reported experiments.

Rather than evaluating sine and cosine at every sample, each harmonic is generated by a two-state rotation:

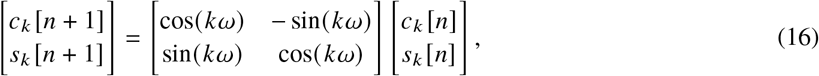

where *ω* = 2*π f*_PLI_ / *f*_*s*_. The states are periodically renormalized to limit floating-point drift. When the tracker supplies a new *f*_PLI_, only the rotation coefficients are changed; the oscillator phase remains continuous.

### 2.5 Supervisory mains-frequency tracker

The frequency tracker observes the band-pass output before the static notch. At *f*_*s*_ = 2 kHz it stores a one-second, 2000-sample circular buffer and evaluates an estimate every 250 ms. The mean is removed, a Hann window is applied, and a zero-padded real FFT with *N*_FFT_ = 8192 is computed. Within the search interval [48, 52[ Hz, the largest spectral peak is refined by three-point parabolic interpolation in log power. The confidence statistic is the peak-to-median power ratio within the search interval, and an estimate is accepted only when this ratio exceeds 2.5.

Let 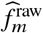 denote an accepted estimate at supervisory update *m*, and let *f*_*m*_ denote the oscillator frequency before that update. The implementation uses smoothing coefficient *β* = 0.5 and bounds each accepted update to Δ *f*_max_ = 0.25 Hz:

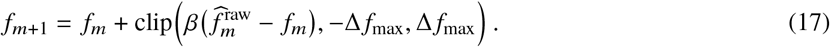

The raw estimate is additionally clipped to the search interval. If the confidence condition is not met, no update is issued and the oscillator retains its last accepted frequency. When an update is accepted, only the oscillator’s rotation coefficients change; its phase states remain continuous.

This two-rate architecture separates the functions of the adaptive stages: the NLMS weights estimate harmonic amplitude and phase, whereas the supervisory estimator tracks slowly varying frequency. The tracker introduces a one-window startup requirement and a periodic FFT task, so its timing and memory are reported separately from the sample-by-sample fast path.

### 2.6 Implementation accounting

The SciPy call `butter(N=4, btype=“bandpass”, output=“sos”)` produces four SOS and therefore an eighth-order band-pass. In direct-form-II transposed form, one SOS uses five multiplications, four additions/subtractions, and two dynamic state values. The fast-path account in table 2 assumes two harmonics, zero leakage, precomputed *μ*/*K*, and recursive oscillator coefficients.

**Table 1.**
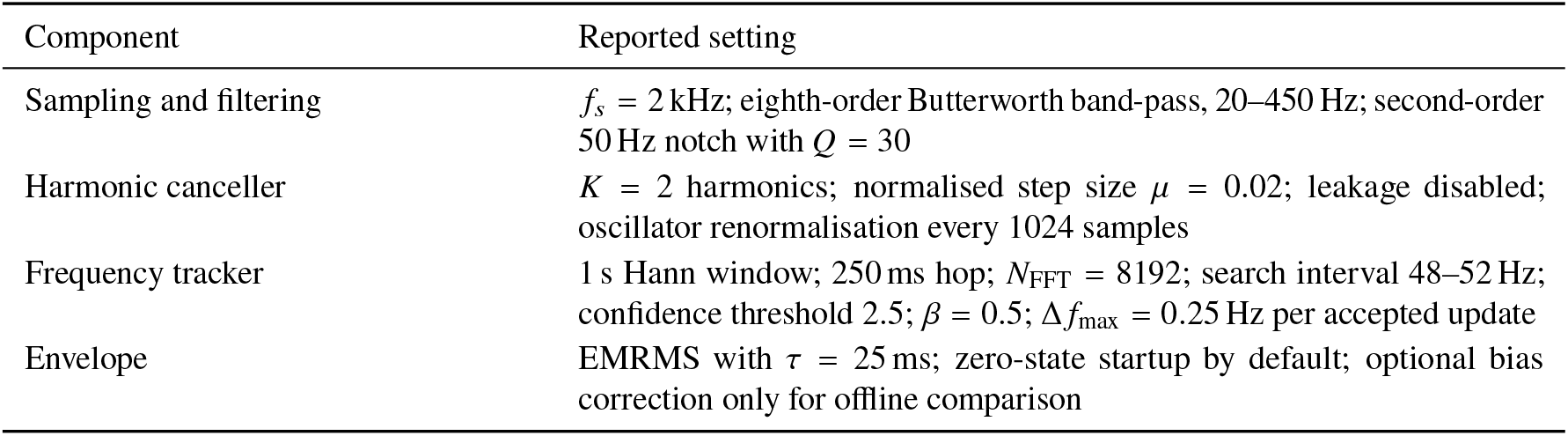
Default parameters of the reported digital implementation.

**Table 2.**
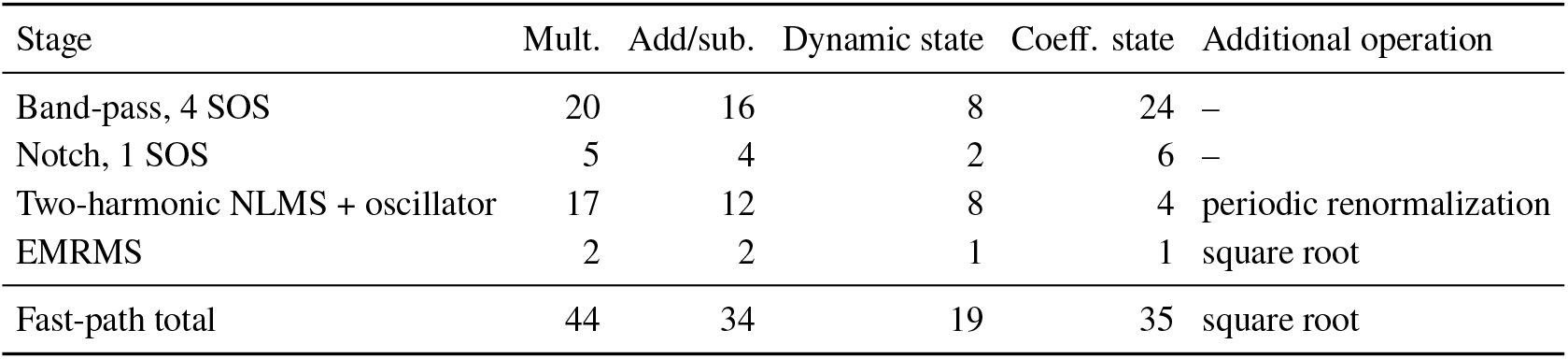
Per-sample fast-path arithmetic and dynamic state for the implemented one-channel chain. The periodic tracker is listed separately because its FFT is not a per-sample fast-path operation.

| Stage | Mult. | Add/sub. | Dynamic state | Coeff. state | Additional operation |
| --- | --- | --- | --- | --- | --- |
| Band-pass, 4 SOS | 20 | 16 | 8 | 24 | – |
| Notch, 1 SOS | 5 | 4 | 2 | 6 | – |
| Two-harmonic NLMS + oscillator | 17 | 12 | 8 | 4 | periodic renormalization |
| EMRMS | 2 | 2 | 1 | 1 | square root |
| Fast-path total | 44 | 34 | 19 | 35 | square root |

The supervisory tracker stores a 2000-sample circular buffer, a window vector, FFT workspace, and search metadata. Its memory and execution cost should be budgeted in a lower-rate task rather than hidden in the fast-path table.

### 2.7 Synthetic signal construction and evaluation protocols

All numerical performance results use controlled synthetic signals so that the line frequency, harmonics, clean reference, and activity windows are known. For generated channel *c*, the contaminated input has the general form

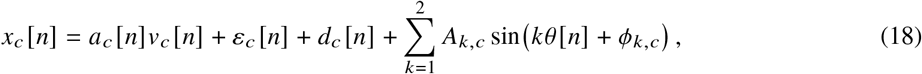

where *v*_*c*_ [*n*] is unit-RMS Gaussian noise passed through the same 20–450 Hz band-pass, *a*_*c*_ [*n*] is a smooth activity envelope formed from hyperbolic-tangent onset and offset transitions with a 30 ms edge scale, *ε*_*c*_ [*n*] is broadband background noise, *d*_*c*_ [*n*] is an optional low-frequency drift, and

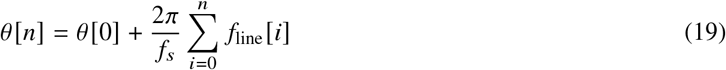

defines the drifting line phase. Deterministic random seeds are specified in the figure-generation script and recorded with the release.

#### Drifting-line experiment

A 10 s record uses seed 9, three generated activity intervals, a 3 mV fundamental, a 0.42 mV second harmonic, and a line-frequency trajectory varying between approximately 49.86 Hz and 50.52 Hz. Static notch, fixed-frequency harmonic NLMS, and tracked harmonic NLMS are compared against the clean signal processed by the band-pass alone. The first 1.2 s is excluded from reconstruction metrics to remove tracker and filter startup.

#### Twelve-channel ablation

Twelve generated channels use seed 21 and are sampled at 2 kHz for 6 s. Each channel has independently generated activity timing, amplitude, background noise, and phase offset, while all channels share the controlled drifting PLI source. A 0.8 Hz, 0.08 mV drift term is added with a channel-dependent phase. The first 2 s is excluded from reconstruction metrics. These channels are repeated software cases, not independent participants.

#### Timing benchmark

The host-Python implementation processes 30 000 samples. Total synchronous time, fast-path time excluding the tracker, and accepted tracker-update time are recorded separately. These values are reference-environment measurements and are not interpreted as MCU worst-case execution time, RTOS jitter, power, or battery performance.

The release also includes `scripts/validate_ninapro.py`. It requires explicit local MAT files and a user-specified scale-to-volts factor, exits with an error when recordings are absent, and never substitutes generated data. The command injects controlled PLI into each clean channel, produces channel-level metrics, and aggregates results by subject.

### 2.8 Evaluation metrics

For a clean band-pass reference *s*_ref_ [*n*] and a conditioned estimate 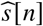, reconstruction SNR over an explicitly stated index set ℐ is

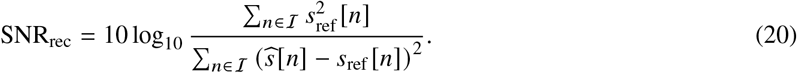

The tracker RMSE is computed at accepted update times after the one-window startup:

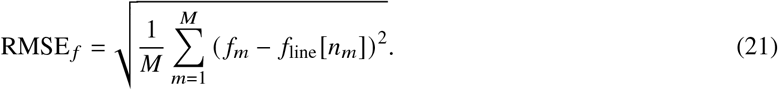

Envelope alignment searches integer lags within ±150 samples and selects the lag maximising Pearson correlation. A positive lag means that the estimate is delayed relative to the reference. After alignment, normalised RMSE is defined as

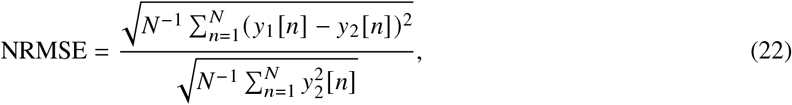

where *y*_2_ is the running-RMS reference. Whole-record reconstruction SNR includes both activity and rest after startup; activity-only SNR and rest-envelope summaries are therefore also reported for the drifting-line experiment.

## 3 Results

### 3.1 CMRR sensitivity screen

Figure 1 evaluates equation (4) with *Z*_in_ = 10 GΩ, CMRR_INA_ = 110 dB, Δ*Z* ∈ [10^2^, 10^6^ Ω], and three assumed DRL attenuation values. The curve identifies sensitivity to impedance mismatch but is neither a measured CMRR curve nor evidence of loop stability or electrical safety.

**Figure 1.**
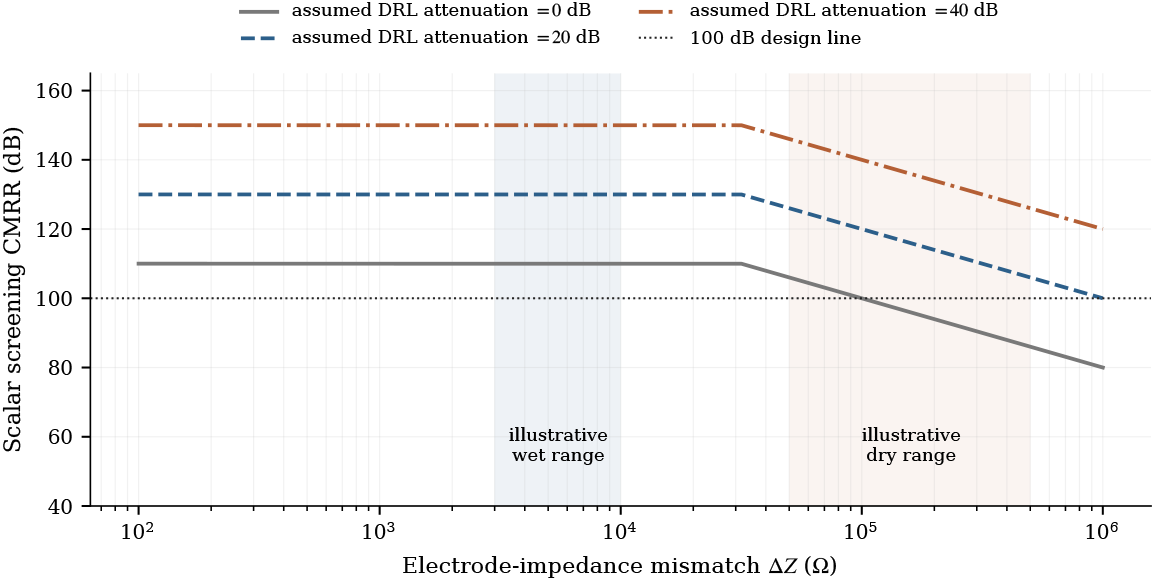
Scalar CMRR sensitivity screen. The assumed DRL term is closed-loop attenuation at the frequency of interest. The shaded impedance ranges are illustrative. Complex electrode impedance, phase interaction, cable parasitics, component tolerances, saturation, safety-current constraints, and loop stability are omitted.

### 3.2 Envelope trade-off and startup behaviour

Figure 2 compares running-sum RMS with EMRMS on the same generated bursts. The *L* = 50 estimator responds quickly but is more variable; the *L* = 200 estimator is smoother and retains a longer finite history. EMRMS follows an intermediate smoothness–response trade-off using one power state. The onset panel also distinguishes zero-state EMRMS from optional startup bias correction.

**Figure 2.**
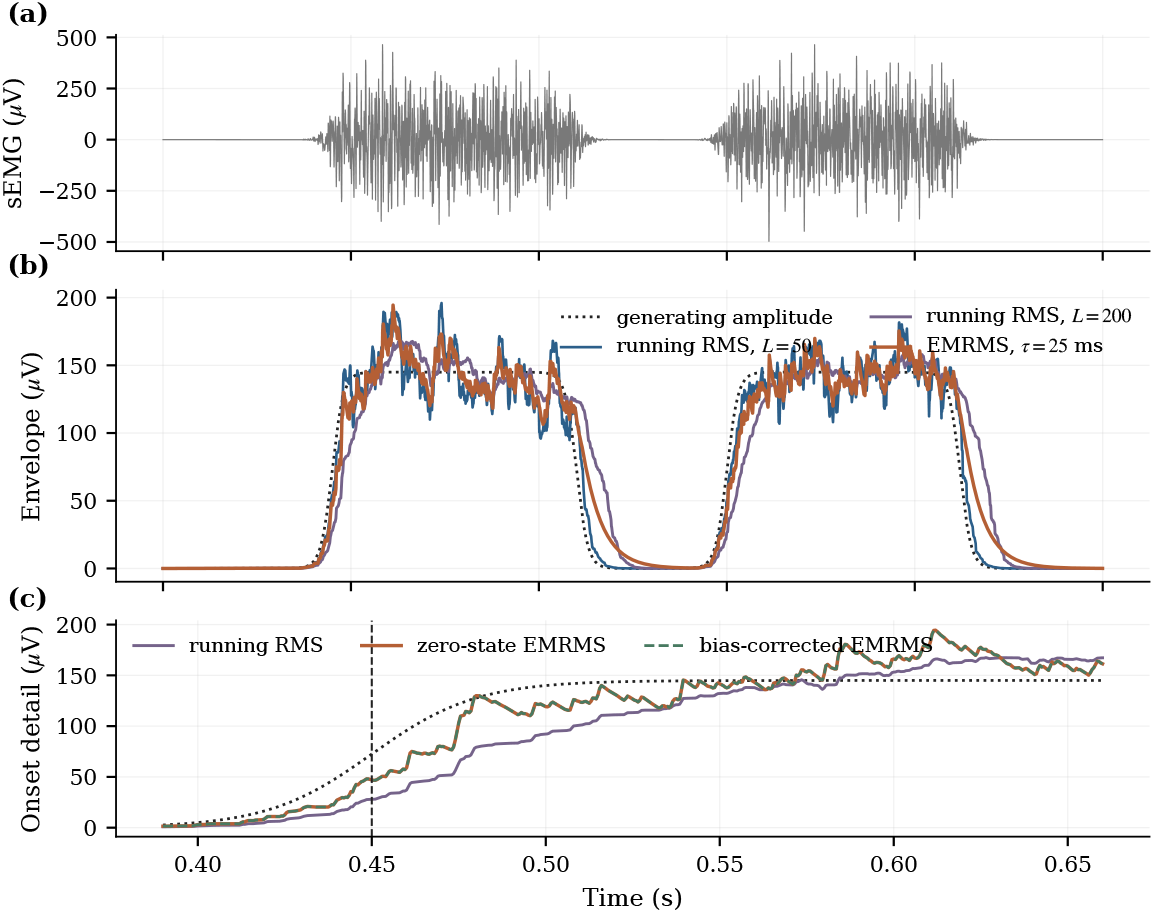
Envelope-estimator comparison. (a) Generated band-limited sEMG-like bursts. (b) Generating amplitude, running-sum RMS estimators, and EMRMS. (c) Onset detail, including optional startup bias correction. All estimators are causal.

**Figure 3.**
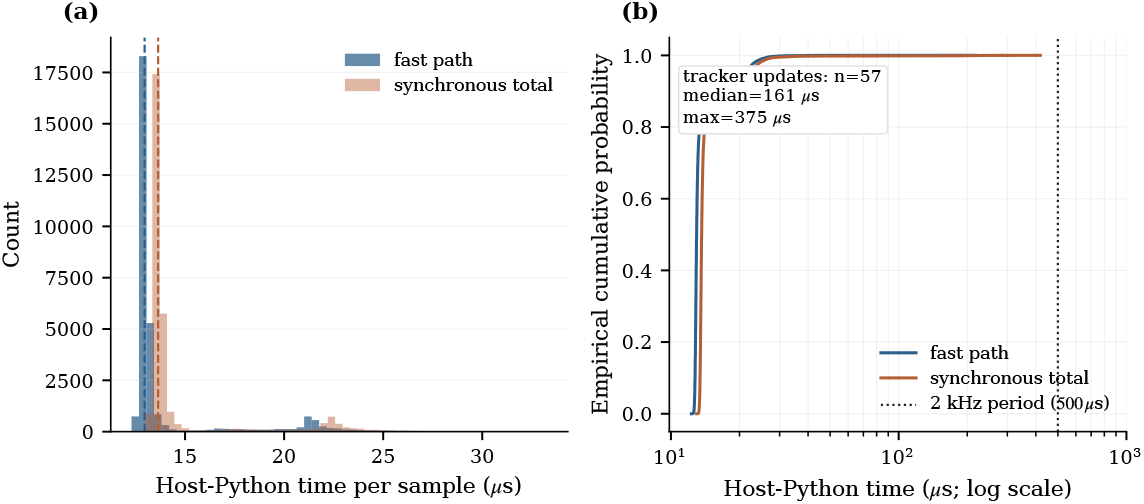
Host-Python timing. (a) Fast-path and synchronous-total distributions, truncated at the 99.7th percentile for readability. Empirical cumulative distributions and the 500 µs sampling period. The inset reports accepted tracker-update cost.

**Figure 4.**
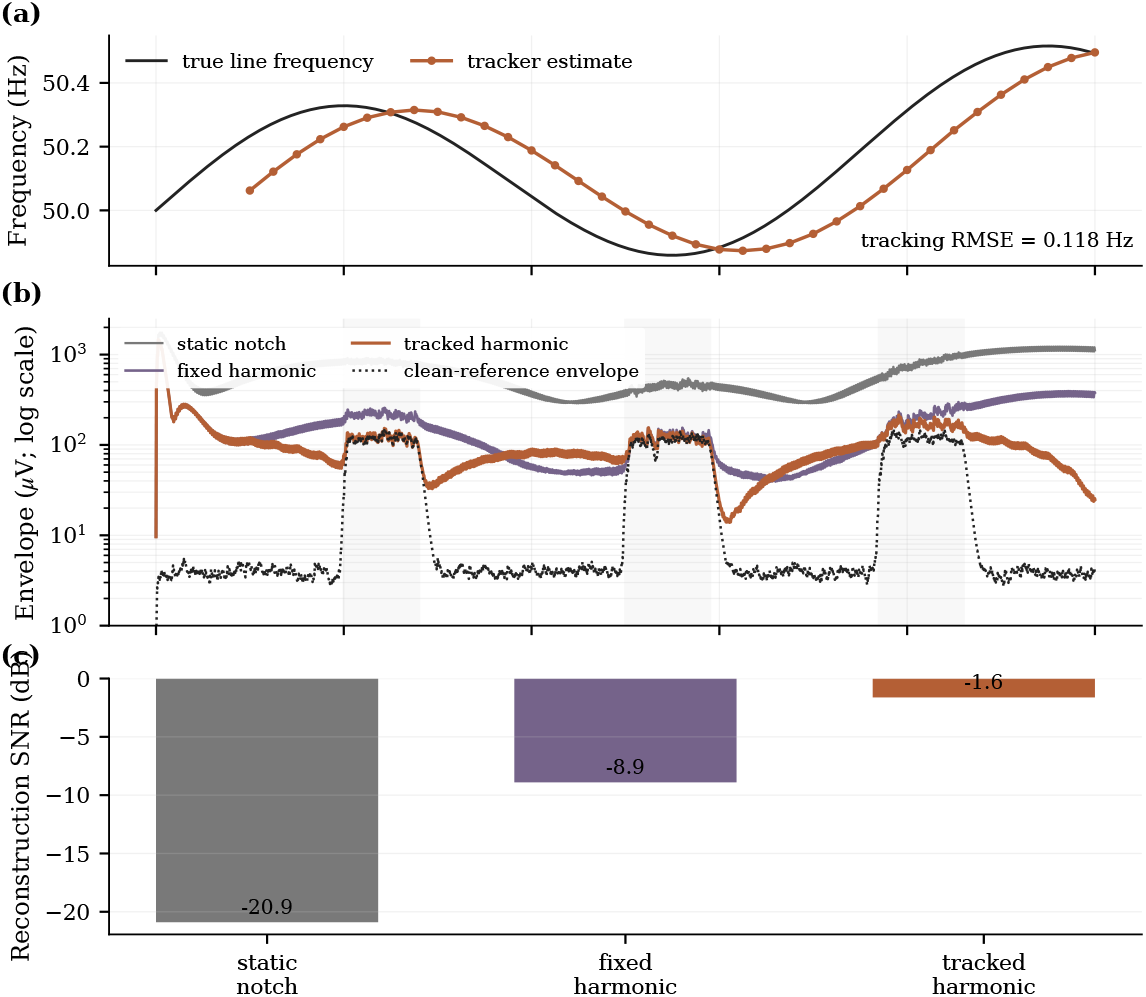
Controlled frequency-drift experiment. (a) True line frequency and rolling tracker estimate. (b) EMRMS envelopes after a static notch, a fixed-frequency two-harmonic NLMS, and the tracked two-harmonic NLMS, with the clean-reference envelope shown as a dotted line on a logarithmic amplitude scale. (c) Reconstruction SNR against the clean band-pass reference.

### 3.3 Reference execution time

The host-Python fast path had a median of 13.0 µs and a 99th percentile of 24.9 µs. Total synchronous time, including tracker pushes and periodic FFT updates, had a median of 13.6 µs and a 99th percentile of 26.6 µs. Accepted tracker updates required a median of 161 µs; the measured total maximum exceeded the 500 µs sample period because an FFT update occurred in the synchronous Python call. This observation supports a two-rate implementation in which the tracker is scheduled outside the hard sample interrupt.

### 3.4 Frequency tracking and correction under drift

In the drifting-line experiment, the tracker achieved frequency RMSE 0.118 Hz after its startup window. The tracked harmonic method improved reconstruction SNR by 19.3 dB relative to the static-notch baseline and by 7.3 dB relative to the fixed-frequency harmonic method. Because whole-record SNR includes long rest intervals with very low reference power, its absolute value remains conservative. Restricting the metric to generated activity intervals gives 3.5 dB for the tracked method, while its median rest envelope is 74.6 µV. These values remain properties of a controlled synthetic ablation rather than clinical signal-quality claims.

### 3.5 Pipeline behaviour

Figure 5 shows one generated record after each processing stage. The first second is marked as tracker warm-up. This interval is a deployment protocol requirement arising from the frequency-estimation window; it is not hidden inside the reported envelope delay.

**Figure 5.**
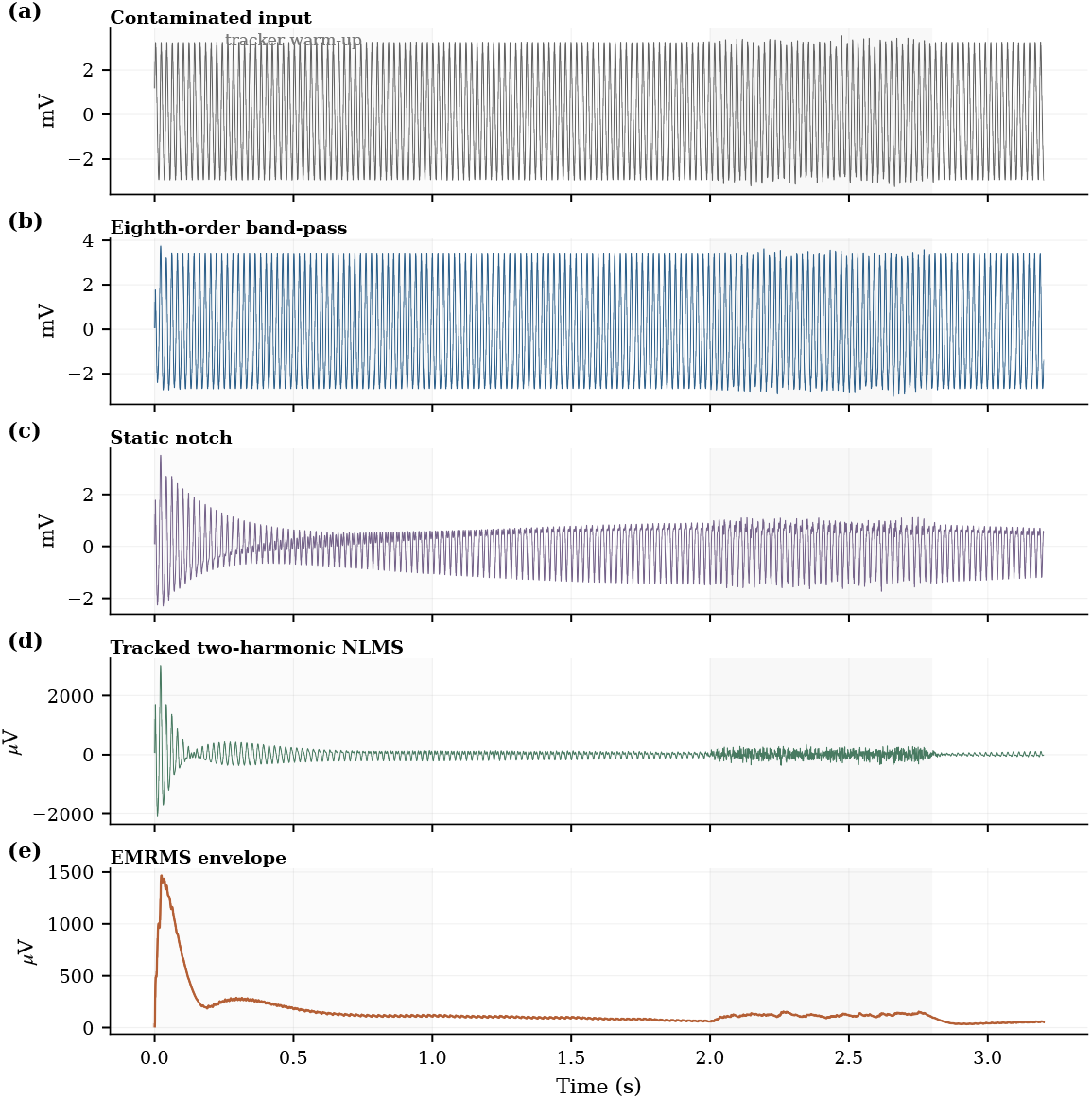
Generated signal through the complete chain: contaminated input, eighth-order band-pass, static notch, frequency-tracked two-harmonic NLMS, and EMRMS envelope. Light grey shading marks tracker warm-up and generated activity intervals.

**Figure 6.**
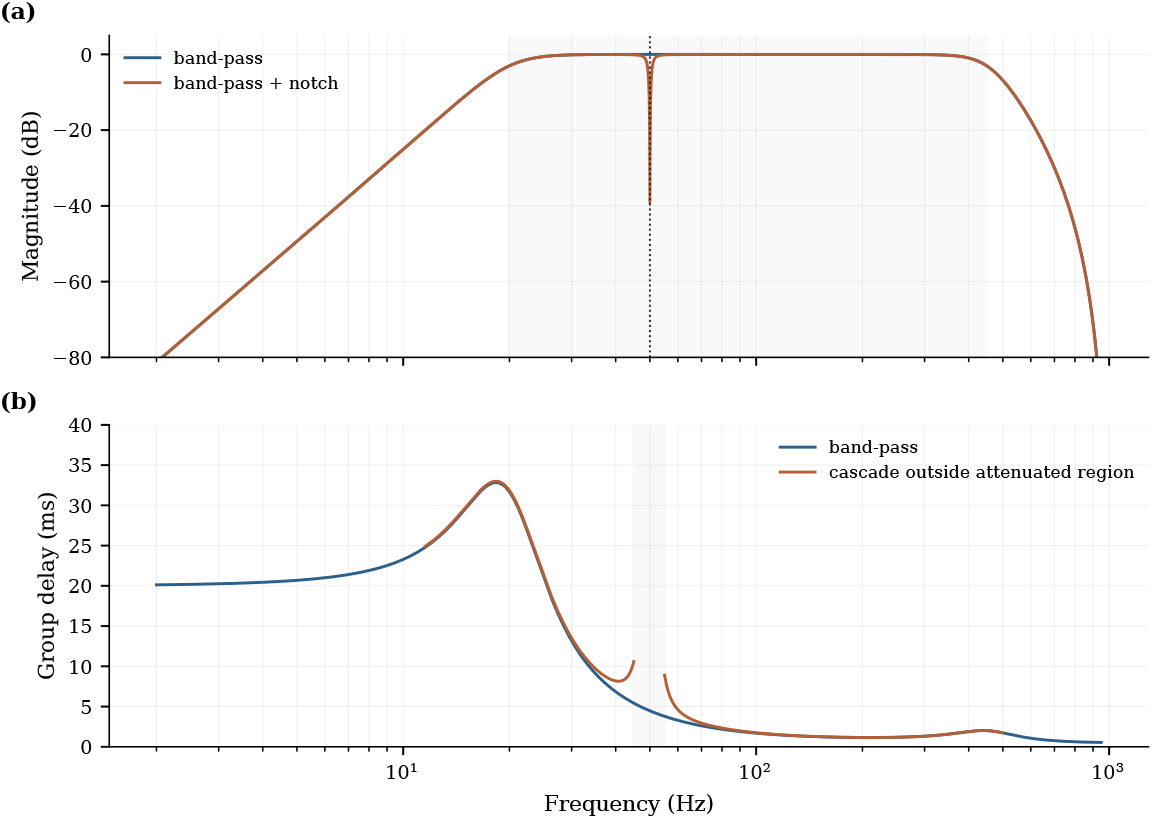
Digital filter response. (a) Magnitude response of the eighth-order band-pass and band-pass-plus-notch cascade. (b) Numerical group delay. The cascade trace is omitted where attenuation exceeds 20 dB and within 45–55 Hz, because phase derivative in a deeply rejected region is not a meaningful signal-delay summary.

**Figure 7.**
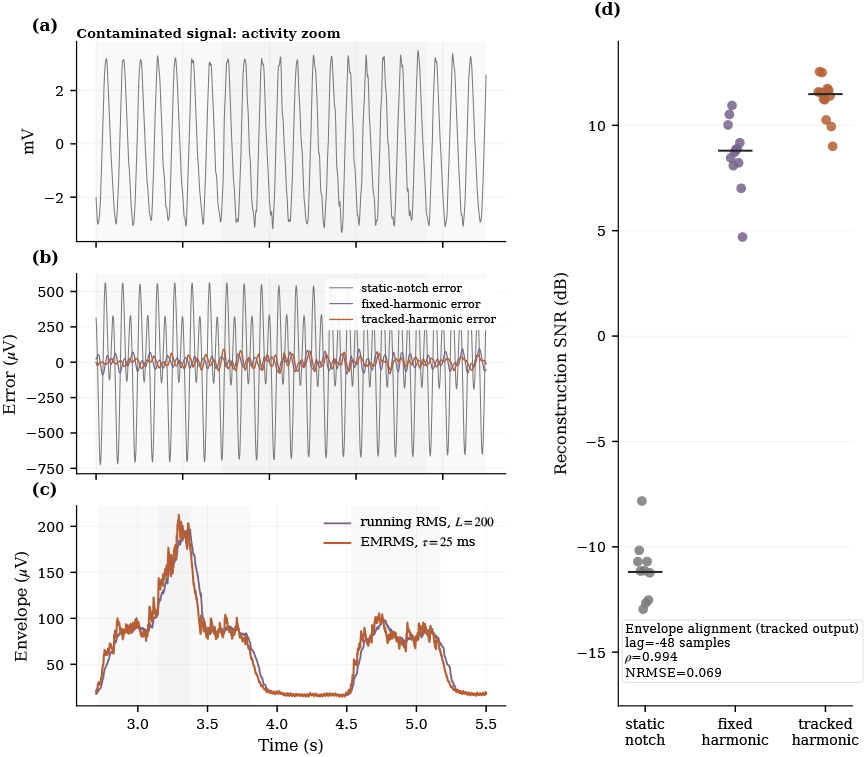
Controlled 12-channel synthetic ablation. (a) Short activity-window zoom of a representative contaminated channel. (b) Reconstruction errors relative to the clean band-pass reference for the three methods over the same zoom. (c) Running RMS and EMRMS envelopes over a wider activity interval. (d) Channel-wise reconstruction SNR; horizontal ticks mark medians. These are generated channels, not Ninapro recordings or participants.

### 3.6 Filter response and group delay

The digital filter delay is strongly frequency dependent. The band-pass contributes approximately 31.7 ms at 20 Hz, 4.5 ms at 50 Hz, 1.7 ms at 100 Hz, and 1.1 ms at 200 Hz. A single constant band-pass delay is therefore only a mid-band summary. Delay close to the deep notch is not interpreted as a useful passband quantity.

### 3.7 Twelve-channel synthetic ablation

Across the 12 generated channels, the median reconstruction SNR was 11.5 dB for tracked harmonic cancellation, 8.8 dB for fixed-frequency harmonic cancellation, and −11.2 dB for the static-notch baseline. The delay-aligned tracked-output EMRMS and length-200 running RMS envelopes had correlation *ρ* = 0.994 and normalized RMSE 0.069. These are descriptive channel-level quantities; no inferential test is reported because the channels are generated software cases rather than independent participants.

### 3.8 Parameter sensitivity

Figure 8 shows the expected trade-offs. Increasing *μ* accelerates cancellation but can raise steady-state adaptation noise and increase the risk of removing genuine narrow-band sEMG components. Shorter tracker windows follow drift more closely in the generated case but offer less spectral selectivity. Cancelling two harmonics materially reduces the residual around the second harmonic compared with a fundamental-only model.

**Figure 8.**
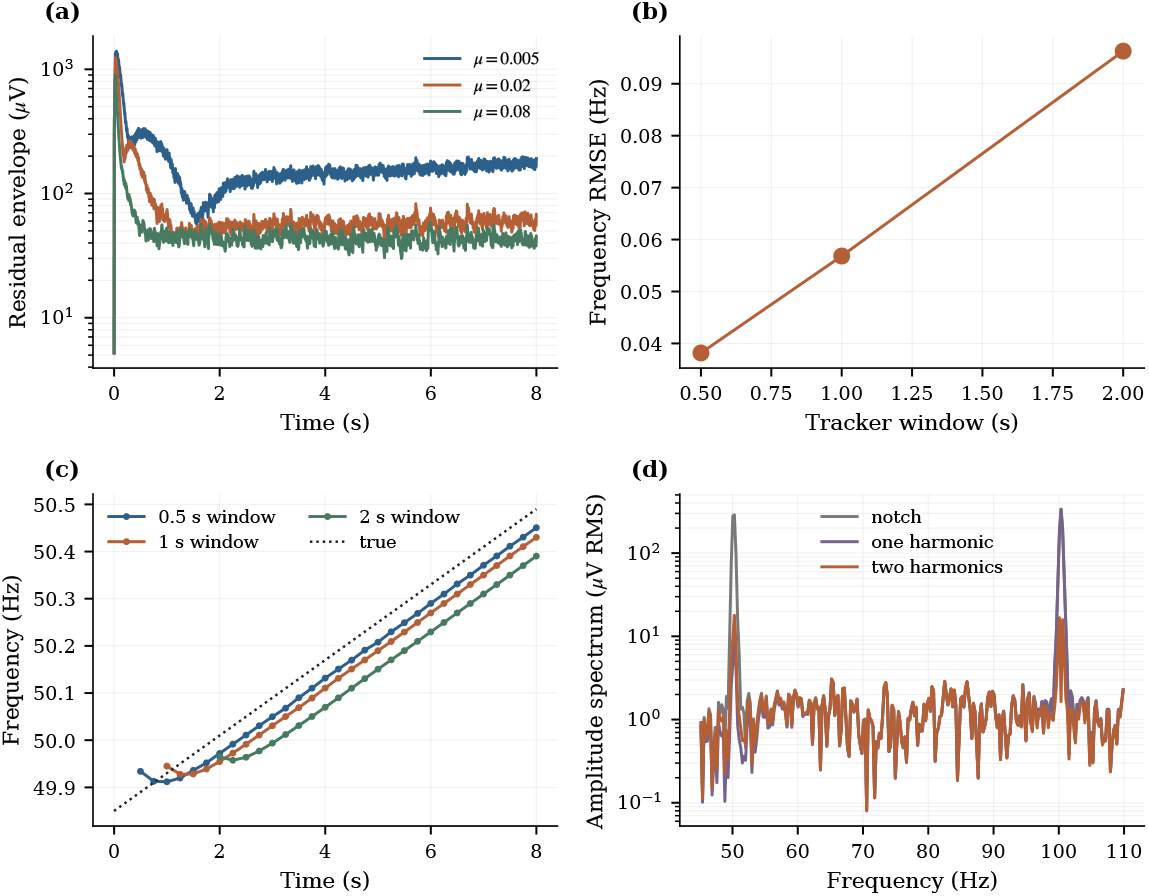
Synthetic parameter sensitivity. (a) Residual envelopes for three NLMS step sizes. (b) Frequency-tracking RMSE for three analysis-window durations. (c) Frequency estimates over time. (d) Spectral comparison of notch-only, one-harmonic, and two-harmonic cancellation.

## 4 Discussion

The results clarify the principal trade-offs in the proposed architecture. The memory advantage of EMRMS is retained when it is compared with an efficient running-sum rectangular RMS: both require constant arithmetic per sample, but EMRMS replaces the *L*-sample history with a single infinite-memory power state. This substitution changes the weighting kernel rather than merely accelerating the same estimator. Centroid delay, onset behaviour, and equivalent window length are consequently more informative than treating *τ* as a universal group delay.

The digital-filter delay is likewise frequency dependent. Near the lower band edge, the eighth-order band-pass contributes substantially more delay than in the mid-band, while delay at the notch centre is not interpretable as a passband timing quantity. Filter group delay, nonlinear envelope timing, tracker warm-up, and software execution time therefore describe distinct phenomena and should not be added into a single universal latency number.

The PLI experiments demonstrate the importance of separating frequency estimation from adaptive amplitude– phase cancellation. Fixed-frequency harmonic NLMS improves substantially over a static notch, but its performance degrades as the line frequency moves away from the nominal value. The supervisory tracker restores much of this loss while keeping the sample-by-sample path free of FFT operations and per-sample trigonometric evaluations. The occasional synchronous timing spike confirms that the FFT task should be double-buffered and scheduled outside a hard sample interrupt, with coefficient updates applied atomically.

The static notch remains in the present chain as a conservative first stage that reduces large initial PLI before adaptation. Its necessity is not yet established, however, and it may distort genuine sEMG energy near the line frequency. A future ablation should therefore compare tracked harmonic cancellation with and without the notch, alongside a published adaptive-notch or joint frequency–harmonic estimator such as Keshtkaran and Yang (2014).

The analog and digital components are not yet experimentally co-validated. The scalar CMRR screen is useful for sensitivity exploration, but a defensible front-end claim requires a specified circuit topology, complex electrode models, component tolerances, DRL loop-gain and phase-margin analysis, common-mode headroom, safety-current constraints, and bench measurements. The present contribution is therefore best interpreted as an analog-aware digital conditioning and modelling framework.

Finally, the current numerical results are transparently synthetic. They establish implementation consistency and controlled known-reference behaviour, not biological generalisability, proportional-control performance, gesture-classification accuracy, or clinical usability. The explicit real-data workflow provides a route to those evaluations without creating a surrogate result when recordings are unavailable.

## 5 Limitations and validation priorities

Although the present results establish the feasibility of the proposed processing architecture under controlled conditions, further empirical validation is needed before broader system-level conclusions can be drawn. In particular, the pipeline should be evaluated using genuine multi-subject recordings from Ninapro DB2 or another clearly identified public dataset. Participants, rather than individual channels, should serve as the primary statistical unit, with results reported through subject-level distributions, confidence intervals, effect sizes, and, where multiple comparisons are performed, an appropriate multiplicity correction. Controlled known-reference experiments would provide an additional means of separating interference suppression from signal distortion. Such experiments should introduce 50 Hz and 60 Hz fundamentals, their harmonics, amplitude modulation, phase discontinuities, and representative frequency-drift profiles into otherwise clean recordings. This design would permit direct measurement of reconstruction error and of any attenuation of genuine sEMG energy near the power-line frequencies.

Evaluation should also extend beyond agreement between envelope estimators. Although envelope correlation provides a useful measure of morphological similarity, it does not directly establish improved control performance. More application-oriented outcomes should therefore include contraction-onset timing, false activations during rest, proportional-control error, and gesture-classification accuracy. These measures would clarify whether the reductions in interference, memory use, and envelope delay translate into practically meaningful benefits for myoelectric interfaces.

The analog analysis presented here likewise remains a pre-prototype model rather than a verified hardware implementation. Circuit-level SPICE analysis and common-mode bench injection are required to compare the measured frequency-dependent CMRR_eff_ (Ω) with the prediction of equation (1). Such validation should account for DRL loop stability and phase margin, common-mode input and output headroom, protection-network effects, electrode-impedance variation, and patient-safety current limits. Until these measurements are available, the CMRR formulation should be interpreted as a design and sensitivity analysis rather than as evidence of realised analog performance.

The embedded properties of the two-rate architecture also require measurement on a specific microcontroller platform. A hardware port should report cycles per sample, worst-case and percentile execution times under realistic interrupts, deadline misses, RAM and flash usage, supervisory-tracker scheduling overhead, and energy per processed sample. A fixed-point implementation would additionally allow direct comparison between floating-point output and Q31 or Q15 arithmetic, including quantisation error, saturation behaviour, and numerical stability.

Finally, the robustness of the supervisory frequency tracker has not yet been characterised across all relevant operating conditions. Particular attention should be given to weak-PLI intervals, strong voluntary sEMG components near 50 Hz, abrupt changes in mains frequency, and failures of the confidence-gating mechanism. A practical implementation should include a conservative fallback policy that freezes or gradually resets the oscillator frequency whenever the spectral estimate is insufficiently reliable, thereby preventing an unstable or erroneous estimate from degrading the adaptive canceller.

## 6 Conclusion

This study presents a reproducible low-latency sEMG conditioning framework that separates common-mode sensitivity analysis, digital filtering, mains-frequency estimation, harmonic amplitude–phase cancellation, and envelope extraction. The rectangular-RMS baseline is implemented with a running sum, the Butterworth band-pass is correctly identified as eighth order, and EMRMS is characterised by one-state memory, centroid delay, equivalent window length, and an explicitly indexed optional startup correction. Frequency drift is handled by a confidence-gated supervisory estimator with bounded updates rather than being attributed to the NLMS weights, while a recursive oscillator avoids per-sample trigonometric evaluation. Controlled synthetic experiments show clear improvements for tracked harmonic cancellation over fixed-frequency and static-notch baselines, and host timing supports a two-rate implementation in which the FFT tracker is scheduled separately from the sample fast path. The accompanying code, tests, machine-readable results, and strict real-data command provide a stronger foundation for the remaining multi-subject, analog, and embedded validation.

## Ethics statement

This study reports no newly collected human-participant data. All numerical performance results in the present version are generated signals.

## Acknowledgements

The authors thank the Ninapro consortium for making its datasets available to the research community (Atzori et al., 2014). Ninapro recordings were not used in the synthetic results reported in this version. The authors also gratefully acknowledge the support provided by Colégio Diocesano de São José 5^a^, Macau.

